# Embryonic and aglomerular kidney development in the bay pipefish, *Syngnathus leptorhynchus*

**DOI:** 10.1101/2021.08.17.456591

**Authors:** Bianca R. Maters, Emily Stevenson, Peter D. Vize

## Abstract

In this report we describe the embryogenesis of the bay pipefish, *Syngnathus leptorhynchus*, and the organogenesis of its aglomerular kidney. Early development was analyzed via a series of montages and images documenting embryos collected from the brood pouches of pregnant males. Despite differences in terminal morphology between pipefish and common teleost models such as medaka and zebrafish, the embryogenesis of these highly advanced fishes is generally similar to that of other fishes. One of the unique features of these fishes is their utilization of an aglomerular kidney. Histological analysis revealed a single long, unbranched kidney tubule in late embryos. The development and structure of this organ was further investigated by cloning the sodium potassium ATPase alpha subunit, atp1a, from *S. leptorhynchus* and developing whole mount fluorescent *in situ* hybridization protocols for embryos of this species. Fluorescent stereoscopic and confocal visualization techniques were then used to characterize the 3D morphology of aglomerular kidneys in intact embryos. In all embryonic stages characterized, the aglomerular kidney is a single unbranched tube extending from just behind the head to the cloaca.

## Introduction

Pipefish are members of the family Syngnathidae, which also encompasses the seahorses and seadragons. At a higher level, they belong to the super-order Percomorpha, which encompasses the sticklebacks, medaka, puffers, sea bass and snappers, amongst many others. The Syngnathids are highly evolved fishes with a number of unique anatomical, behavioural, biomechanical and physiological features that make them extremely interesting as a developmental system (Ahnesjo et al., 2011). An example of these unique adaptations that is of particular interest is the aglomerular kidney, which as the name implies, lack glomeruli and function by a process different to that of other vertebrate kidneys (Marshall et al. 1930; Smith 1932).

The early vertebrates evolved in fresh water and moved to the oceans (Smith 1953). These origins are reflected by the osmotic imbalance that exists between the bodies of marines fishes and the surrounding sea water, which has over twice the salt of their blood (Beyenbach 2004). In order to avoid being dehydrated by the surrounding environment, marine fishes needed to develop systems to minimize water loss. In fresh water fishes the kidney functions to excrete excess water. The filtration resorption system utilized by their kidneys works very efficiently when water conservation is not an issue, but it is energetically very costly in a dehydrating environment. Different marine fishes have solved this problem in different ways-in sharks and rays by retaining high concentrations of urea in the blood; so high that fresh water flows inwards through their gills (Vize and Smith, 2004). In many marine teleosts this issue has been addressed by reducing glomeruli numbers and enhancing secretion/excretion by the kidney (Smith 1953). In some advanced fishes the glomeruli are lost completely and the kidney functions solely via secretion and excretion across the tubule epithelia. The Syngnathids are the only known group to have completely aglomerular kidneys in both embryos and adults, although other fishes have partially or wholly aglomerular kidneys in adults (Marshall et al. 1930; Beyenbach 2004; Ozaka et al., 2009).

In this report the embryonic development of a species of an aglomerular pipefish, *Syngnathus leptorhynchus* (Girard, 1854), the bay pipefish, is described. Embryos were collected from the brood pouches of pregnant males over a number of seasons in the Pacific Northwest. We also report the cloning of the bay pipefish sodium-potassium ATPase (atp1a) that drives many membrane transport processes, and use this clone to develop fluorescent *in situ* hybridization technology in wholemount pipefish embryos. This technique was then used to visualize the organogenesis of an aglomerular kidney for the first time.

## Materials and Methods

### Specimen collection permits

Permit numbers from Bamfield Marine Sciences Center AUPE were RS-06-21, RS-08-27 and RS-09-05. In 2009 regulations also required a permit from Fisheries and Oceans Canada, and this was XR 103 2009.

### Embryo collection methodology

It is necessary to euthanize pregnant males for embryo removal from the brood pouch as all efforts to remove embryos from anaesthetized males resulted in the animals death within 48 hours. Following euthanization and pithing, an incision is made along the anterior and lateral edges of the pouch. When embryos are in early stages of development (pre-pigmented eyes) the embryos are firmly attached to the surface of the pouch by deposits of extracellular matrix (Kornienko 2001). The entire pouch and the attached embryos can then be removed and transferred to a dish of seawater. Embryos are detached from the pouch by running a blunt probe between the surface of the pouch and the rows of embryos, which will detach in a sheet. Later stage embryos are not attached to the pouch and will remain in the pouch cavity, from where they can be collected by rinsing with seawater. Embryos remain within the fertilization envelop up until close to hatching, but the envelop is very fragile from eye pigmentation stages onwards, and is usually disrupted during embryo removal unless great care is taken. Upon collection embryos were fixed in MEMFA (0.1 M MOPS pH7.4, 2 mM EGTA, 1 mM MgSO_4_, 3.7% formaldehyde) for 2 hours at room temperature, then transferred to 100% methanol for long term storage. For photodocumention, embryos were slowly rehydrated through a series of methanol:tris buffered saline (TBS, 12 mM Tris.HCl pH7.5 150 mM NaCl) at room temperature starting at 98% (methanol:TBS), then 95%, 90% 75%, 50%, 25% and finally straight TBS.

### Histology

For sectioning embryos in 100% methanol were mounted in JB4 plastic sectioning medium (Polysciences cat. no 00226-1). Dehydrated samples were infiltrated with 10 ml JB4 solution A plus catalyst for two days, with two changes of solution per day, with gentle shaking at room temperature. This solution was then mixed with JB4 solution B, placed in a mold, and sealed. After curing overnight, blocks were trimmed and sectioned at a thickness of 2 microns with a diamond knife and microtome. Sections were placed on water droplets and dried onto slides, then stained with hemotoxylin (Sigma GHS-3-32) and eosin (Sigma HT110-2-3) for two minutes each, washed with water, then dehydrated with ethanol and mounted with Permount (Fisher 024578).

### Cloning

Late stage embryos to be used for RNA isolation were homogenized in Trizol (Invitrogen) and transported to the laboratory at room temperature, where they were then frozen at -80°C for storage. RNA was isolated from mid-hatching stage embryos according to the Trizol manufacturer’s protocol and stored in water at -80°C.

Total RNA was converted to cDNA using AMV reverse transcriptase (Promega) using an oligo dT primer using the Promega supplied protocol and reaction buffer. Following incubation at 45°C for one hour the reverse transcription reaction was heated to 85°C for ten minutes then frozen for storage. 2 ul of the cDNA reaction was used in a 50 ul PCR reaction containing 1.5 mM MgSO_4_ and using Pwo DNA polymerase (Fermentas).

Primers for RTPCR were based on conserved amino acids in NaK ATPase from other species and were GCTCACCTTGGATGAGCTTC and GATTCACCGGTCAAAGAGAGGA. The blunt PCR product was ligated into pJET1.2 (Fermentas) and sequenced using a primer against the single RNA polymerase promoter in this vector, T7. The results demonstrated that the clone encoded an atp1a cDNA, but that the insert was in a sense orientation relative to the T7 promoter, and therefore not suitable for generating the antisense RNA probes *in situ* hybridization requires. The full length insert of this plasmid was therefore excised with *Bgl*II and ligated into the *Bam*HI site of pCS108, which has RNA polymerase promoters on both sides of the multiple cloning site. A clone with an insert was selected and the 1,100 bp insert fully sequenced on both strands. The sequence of the plasmid, named pSLatp1a, is available from Genbank (accession number GU735082.1 for DNA, ADE80883.1 for protein).

### Fluorescent in situ hybridization

Standard wholemount fluorescent *in situ* techniques developed for other large embryos were used (Vize et al., 2009,; Lea et al. 2012). Due to the dermal plates and the longer than normal length of late stage embryos, some modifications to published protocols were required. These include extending incubation in proteinase K to enhance probe penetration and in late stage embryos, removal of the head and tail to give sufficient probe access. Antisense RNA probes labeled with DNP were used in all *in situ* hybridizations and probe detection was performed using a HRP-coupled anti-DNP antibody (PerkinElmer) and a fluorescein coupled tyramide (Vize et al., 2009). Samples were photographed using a Leica FLIIIZ fluorescent stereoscope and a Leica HQ:F filter, or on a Leica SL spectral confocal system and 10X or 20X long working distance APO objectives. The confocal stacks were processed using AMIRA software (ThermoFisher) to generate 3D models of labelled structures.

## Results

The bay pipefish, *Syngnathus leptorhynchus*, is relatively common in the eel grass beds of the Pacific coast of North America. In the summer months from May to July the majority of males captured via seine netting are pregnant. *S. leptorhynchus* has a complex sealed pouch, akin to that of *S. floridae* (Ripley et al., 2006). All of the embryos illustrated in this report were removed from the brooding pouch of wild-caught fish captured in June or July. As is obvious from some of the data presented below, the colour of embryos changes dramatically during the course of development in the pouch. Early embryos are an intense orange colour due to yolk pigmentation and the lack of melanocytes of more mature embryos. The intensity of the yolk colour declines over time and the yolk sac decreases in size as it is metabolized. In addition, hatching stage fry are darkly pigmented with many melanocytes. When capturing fish for embryo harvesting, the colour of the embryos can be observed through the translucent brood pouch allowing for rough staging in the field and the collection of only appropriate fish. Approximately 20 pregnant males per season were collected by seine netting (under permit) from the eel grass beds adjacent to the Port Alberni Yacht Club on Fleming Island at 48° 53’ 34N and 125° 7’ 7W. Animals were transported to Bamfield Marine Sciences Centre, British Columbia, Canada, were they were kept in tanks on sea tables.

### Embryonic development of Syngnathus leptorhynchus

The techniques used to collect embryos from the brood pouches of pregnant males are described in the Materials and Methods section. After collection embryos that were to be used for photodocumentation were fixed in formaldehyde then stored in methanol at -20°C until needed. Embryos were slowly rehydrated into tris-buffered saline for imaging. High magnification images of embryos were collected using a Qimaging digital camera then merged into a single montage of each embryonic stage using Adobe Photoshop.

The chorion, or egg envelop, of *S. leptorhynchus* has a rubber-like quality and does not harden like the chorions of many fishes that lay eggs externally. The chorion can be manually removed with fine watchmakers forceps without obvious damage to the embryo. To date we have only captured a small number of fish with very early stage embryos so a complete description of cleavage stages is not possible. In figure 1, panel A illustrates the typical appearance of the few batches of very early embryos that we have obtained to date. The early embryo is bright orange in colour, containing numerous lipid droplets. Within the yolk mass the lipid droplets accumulate dorsally directly under a disc of cells representing the late blastula/early gastrula embryo. In different batches this disc is of different diameters, but to date no discs broader than those shown have been observed. The embryo shown in figure 1A is similar to that published by Silva *et al*. in their description of the development of *Syngnathus abaster* (2006) and is quite similar to the pre-gastrulation stages of other fishes, for example salmon (Boyd et al. 2010) and medaka (Iwamatsu 2004a). The next developmental stage identified were post gastulation stages commonly referred to as segmentation stages (Iwamatsu 2004a), as this period of development is when the embryonic muscle blocks known as somites are segmenting from the paraxial mesoderm. Two different segmentation stages are illustrated in Fig. 1. Panel 1B depicts an early segmentation stage. The neural tube is clearly visible, but no retinas are present. Lipid droplets are common in the yolk sac, as they are in all stages up until early hatching stage. Panel 1C depicts a somewhat older segmentation stage. At this point in development a retina is clearly visible, while the tail bud at the posterior-most point of the main body axis remains attached to the yolk. The next major developmental stage is often referred to as the pharyngula stage as there is a distinct post-anal tail, and is represented by Fig. 1 panels D through H. In this developmental period the tail greatly increases in length from the 1 mm of a segmentation stage to approximately 5 mm by the stage depicted in Fig. 1H. In this period the fin blastemas form and most develop fin rays, the mouth forms, and melanocytes proliferate and spread throughout the dermis of the embryo. The yolk sac remains transparent and contains lipid droplets throughout this window. The pharyngula stages are followed by hatching stages. In this final developmental window the snout extends to form the distinctive structure from which the Syngnathidae derive their name, the yolk sac is resorbed, and the body continues to elongate until it is approximately 2 cm long at the time of birth. The embryo depicted in panel 1K represents an individual shortly before birth. Note that the anal fin, shown in the lower inset of the figure, lacks fin rays at all stages imaged.

**Figure 1.**
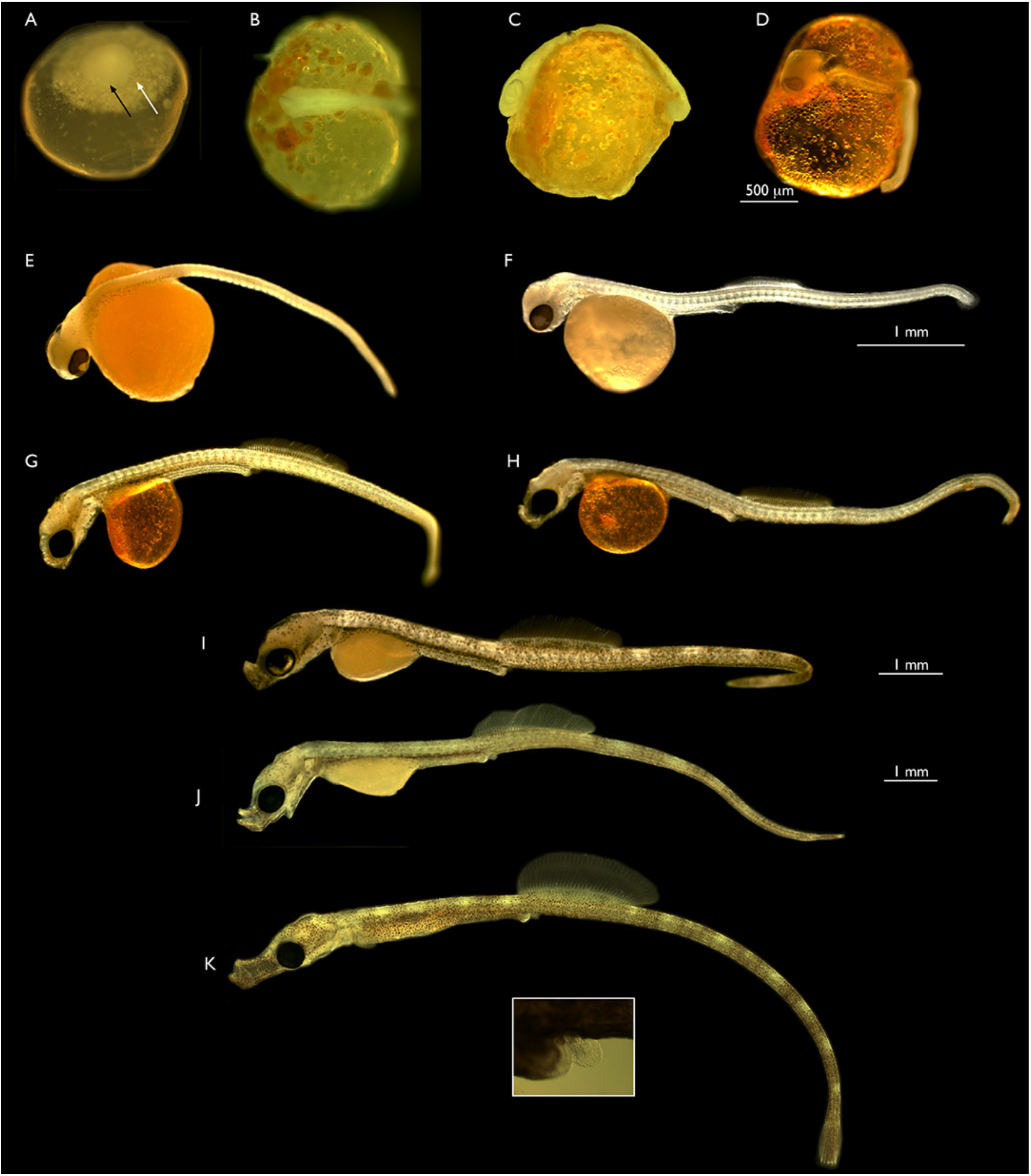
Embryonic development of the bay pipefish. A. Blastula/disc stage. The black arrow indicates the lateral border of the dense disc of cells. The white arrow indicates a larger disc of more dispersed material that the cellular disc sites upon. Approximate equivalent; medaka stage 11-12. B. Early segmentation stage. Anterior end of embryo is to the left. Approximate medaka stage 18-19. C. Late segmentation stage, lateral view. Retina obvious. Approximate medaka stage 22. D. Early pharyngula stage. Retina obvious, extensive somitogenesis. Posterior tail is not attached to yolk sac. Approximate equivalent; medaka stage 25, except the tail is longer and detached in the pipefish. E. Mid pharyngula stage. No snout, no fin blastemas. F. Mid pharyngula stage II. Slight protuberance in future snout region. Dorsal and caudal fin blastemas present, first dorsal fin rays are emerging, no rays in caudal fin. G. Late pharyngula stage. Future snout small. Dorsal fin rays still forming, first rays in caudal fin are apparent. H. Late pharyngula stage. Future snout small. Dorsal fin rays still forming, first rays in caudal fin are apparent. I. Early hatching stage. Snout distinct, jaw visible and mouth can open. Dorsal and caudal fin rays are well developed. Yolk sac is 50% absorbed. J. Mid hatching stage. Snout becoming longer, tail extending, yolk sac remnant rapidly decreasing in size, approximately 25% of original size. K. Late hatching stage, early fry stage, shortly before emergence. Elongated snout, well developed dorsal fin. Yolk sac has been resorbed. Birth imminent. The panel in K illustrates the anal fin in this embryo under higher magnification.

Higher magnification images were also collected to document in greater detail anatomical feature progression during pharyngula, snout and hatching stages of embryonic development (Fig.2). The dorsal and caudal fin buds appear somewhere between the stages depicted in Fig.1E and 1D. There is no mouth present in these stages and other than in fin structures and body length, they look very similar. The fin blastemas first form as translucent aggregations (Fig. 2A) which then increase in size (Fig 2B). At the same time as the mouth is first visible, rays are visible in the dorsal fin blastema only. When the mouth is clearly present, rays have also appeared in the tail fin blastema (Fig. 2D). Hatching stages are characterized by the elongation of the snout, the increase in the size of the dorsal fin, and a major increase in body length (Fig. 2).

**Figure 2.**
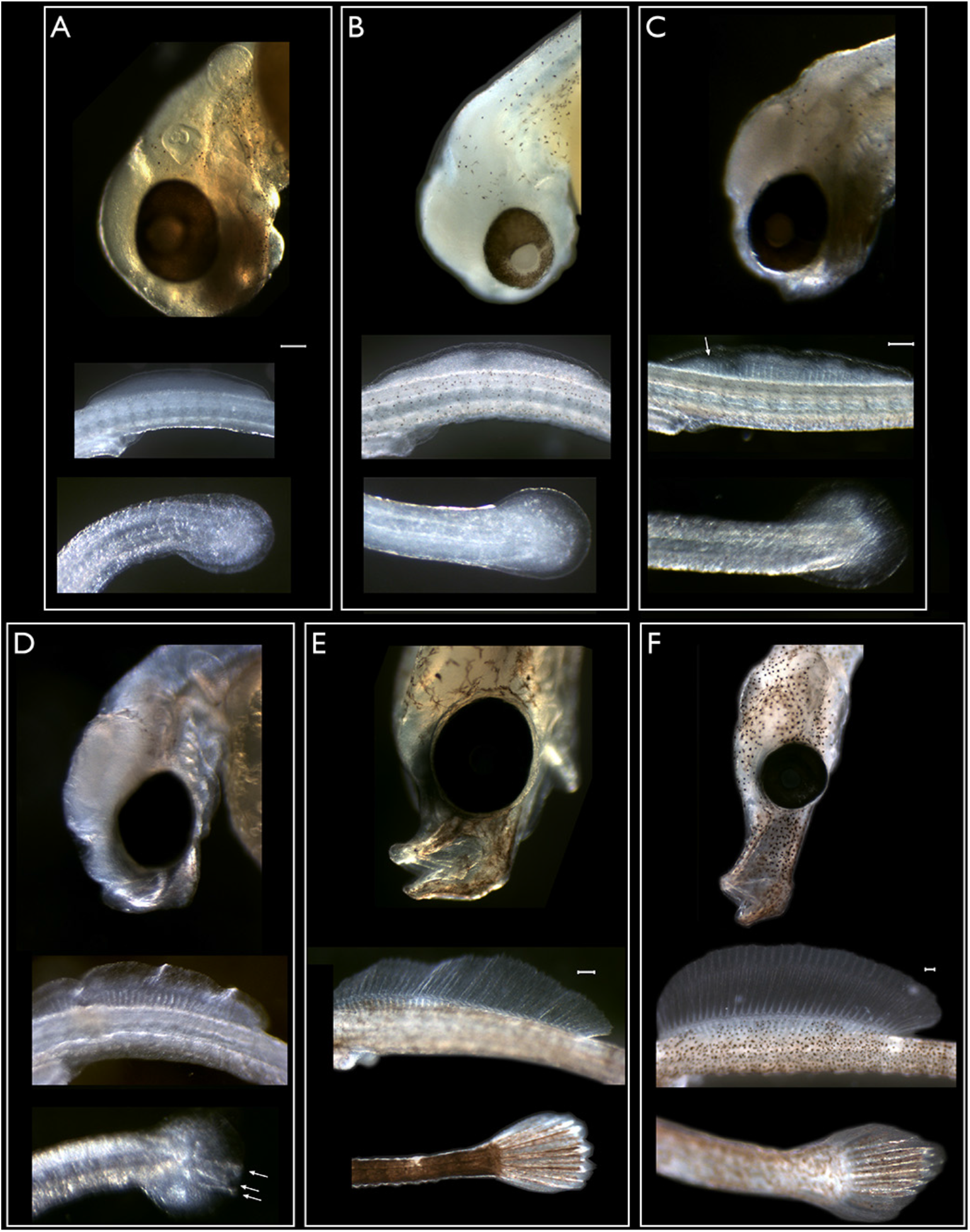
Anatomical features of pharyngula and hatching stage embryos. Scale bars in A, C, E and F represent 100 microns, white arrows indicate fin rays. A. Mid-pharyngula stage, no mouth, dorsal fin and caudal fin blastemas. Otolith visible. B. Similar to A, but slightly older. C. Mid pharyngula stage. Slight protuberance in future mouth region. Dorsal fin rays are emerging, no rays in caudal fin. D. Late pharyngula stage. Future snout small. Dorsal fin rays still forming, first rays in caudal fin are apparent. E. Early hatching stage. Snout distinct, jaw visible and mouth can open. Dorsal and caudal fin rays are well developed. Yolk sac is 50% absorbed. Melanocytes star shaped. F. Late hatching stage, early fry stage, shortly before emergence. Elongated snout, well developed dorsal fin. Yolk sac has been resorbed and birth is imminent.

### Histology

Hatching stage embryos were embedded in JB4 and sectioned to examine the internal anatomy and structure of the aglomerular kidney. In all transverse sections unbranched epithelial tubes corresponding to the nephric system were observed ventral to the large, vacuolated notochord (Fig. 3) and adjacent to the dorsal aorta. In all sections a pair of tubes was observed, each 8 - 10 cells in diameter. No branching and no glomeruli were observed. The kidney tubules look very similar to those described from the greater pipefish, Sygnathus acus (Ozaka et al. 2009).

**Figure 3.**
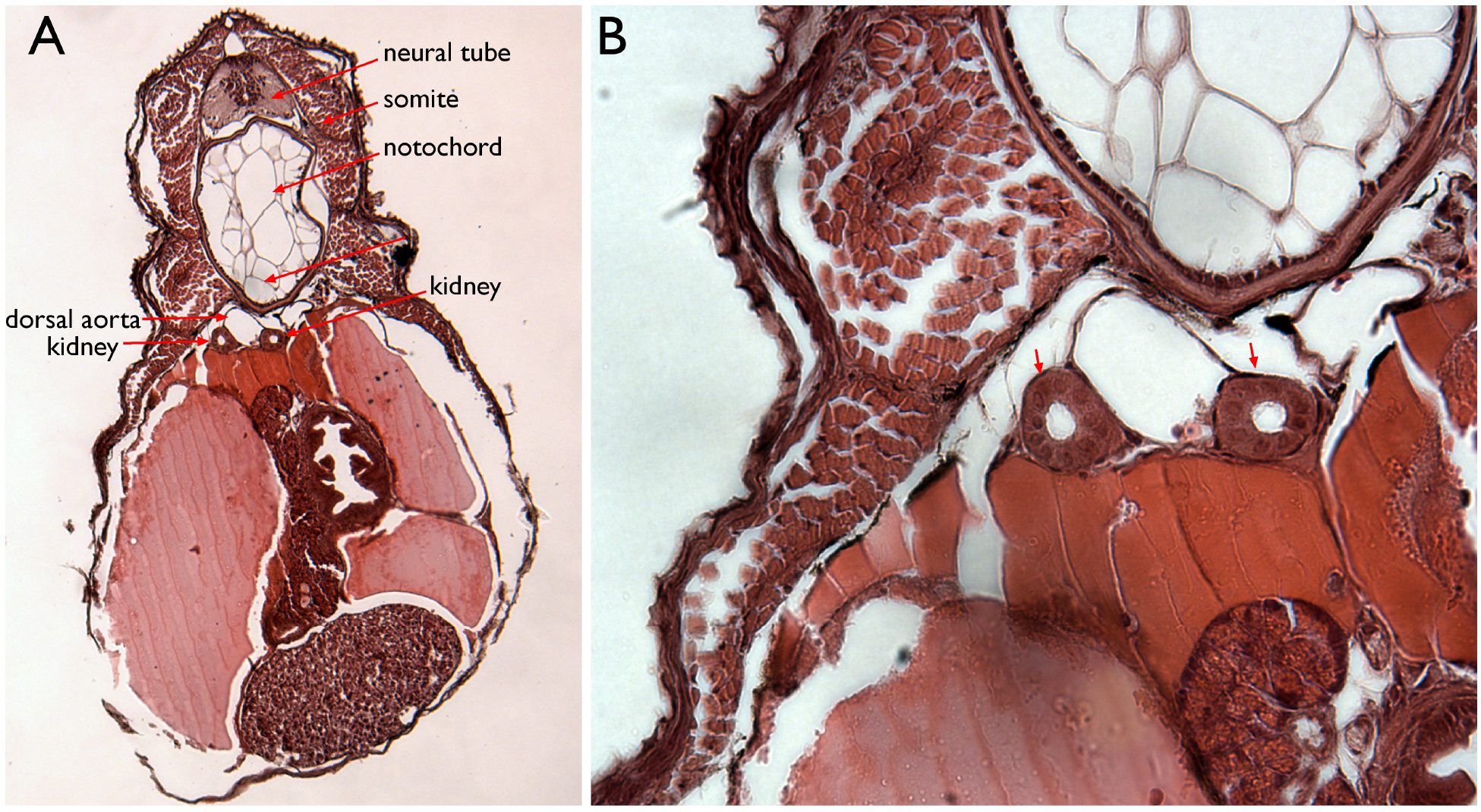
Internal anatomy of mid-hatching stage bay pipefish. A. Transverse section of a JB4 plastic embedded specimen. Various dorsal tissues surrounding the kidney tubules are labelled. B. Enlargement of panel A.

### Cloning the kidney expressed sodium potassium ATPase, atp1a, from S. leptorhynchus

In order to visualize organogenesis of the kidney in this aglomerular species, we cloned a gene that is expressed at high levels in the forming kidney, the sodium-potassium ATPase, atp1a, and used this clone to develop fluorescent *in situ* methodologies for pipefish embryos. The sequence of atp1a1 from zebrafish and Takifugu were used to design PCR primers against a highly conserved protein stretch that was encoded by low redundancy amino acids. These were KDMDDL and HMWFDN/SQ. Redundant oligonucleotides encoding these domains were used to PCR cDNA from late hatching stage embryos. The 1,100 bp PCR product was cloned into pCS108 for further analysis. Sequence of the insert followed by alignment to the NCBI NR protein database revealed that the cloned fragment was most closely related to the atp1a1 subunit of the seabream, followed by the same protein in Tilapia. This sequence was submitted to Genbank (accession number GU735082.1 for DNA, ADE80883.1 for protein) and the clone was named pSLatp1a and used in all subsequent experiments. The predicted protein sequence of the *S. leptorhynchus* protein is highly similar to that of other fish and non-fish atp1 alpha subunits, with long stretches of 100% conservation (Fig. 3).

An antisense RNA probe was generated from pSLatp1a by *Hind*III digestion and transcription with T7 RNA polymerase incorporating DNP labelled ribonucleotides, as described by Vize (2009). The protocol used to perform fluorescent wholemount in situ hybridization (FISH) against pipefish embryos was based on that previously optimized for frog embryos. Some modifications to this protocol were required including extending the period of time in which fixed embryos were treated with protease K prior to hybridization with the DNP-labelled antisense RNA probe. The optimized times for proteinase K permeabilization varied with embryonic stage and other protocol modifications, are described in supplemental table 1. In late pharyngula and hatching stage embryos it was also necessary to remove the head and tail of the embryo to allow reagent access to the interior of these 1 to 2 cm long embryos. Details are once again provided in supplemental table 1. Controls were performed by generating a sense RNA probe. While antisense probes generated a clear signal in the structures identified as kidney tubules in the histology experiments described above, sense probes did not (Fig. 5). Antisense probes also labelled the gut and in some samples, dispersed cells in the skin. The later is expected, as in other fish species this gene is expressed in both the kidney and in a dispersed population of skin cells, possibly ionocytes (Sprague et al. 2001). The staining in the gut is somewhat variable between samples and increases in strength as embryos age. In hatching stage embryos it was often necessary to dissect out the gut to allow imaging of the kidneys.

**Figure 4.**
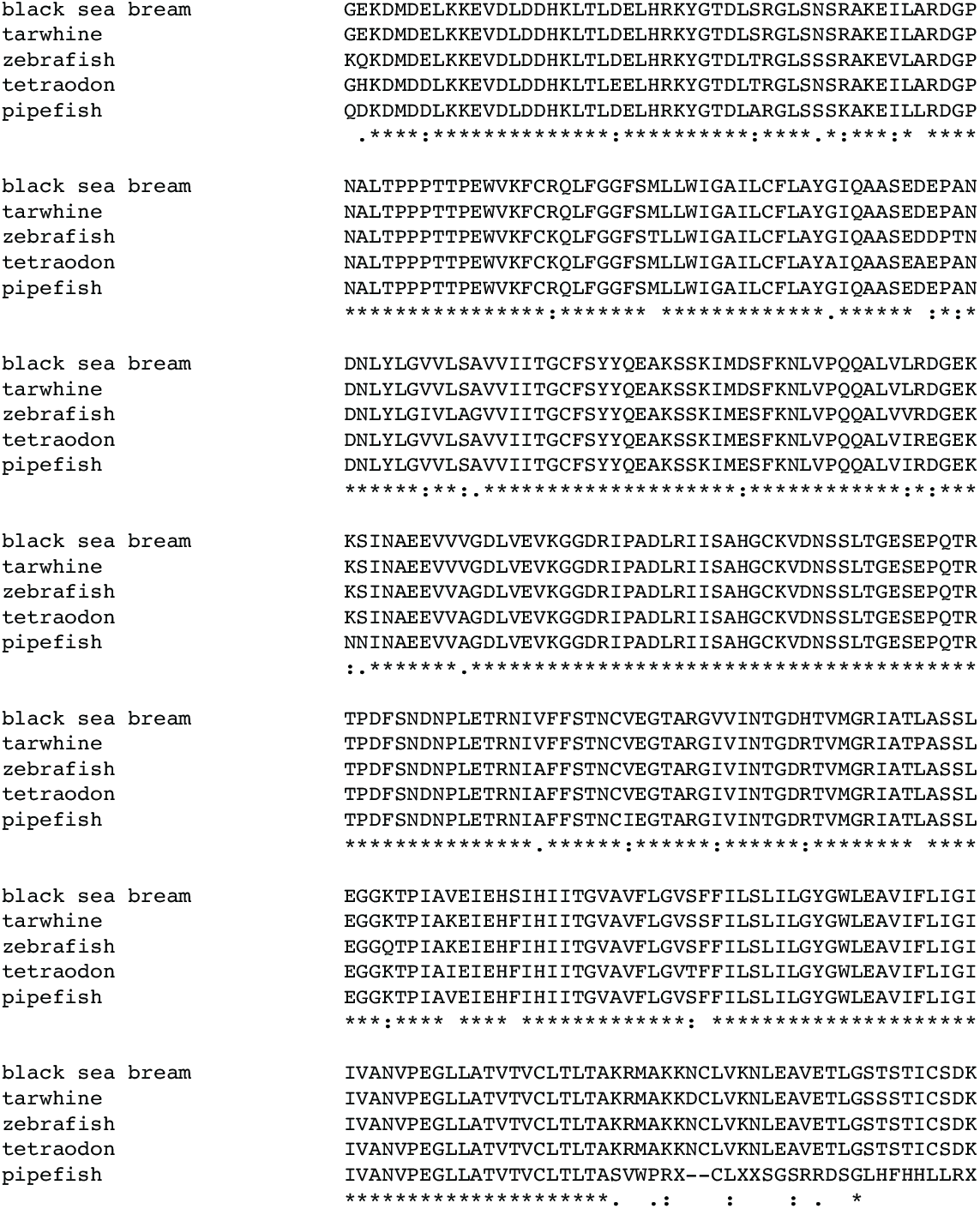
Protein sequence of bay pipefish atp1a aligned to related proteins from various fresh and salt water fishes. Amino acid residues conserved in all species are indicated with a ‘*’, highly similar amino acids with a ‘:’ and similar amino acids with a ‘.’. The NCBI accession numbers of aligned proteins are; black sea bream, ABR10300.1, tarwhine, AAT48993.1; zebrafish AAI63629.1; tetraodon, CAF97804.1.

**Figure 5.**
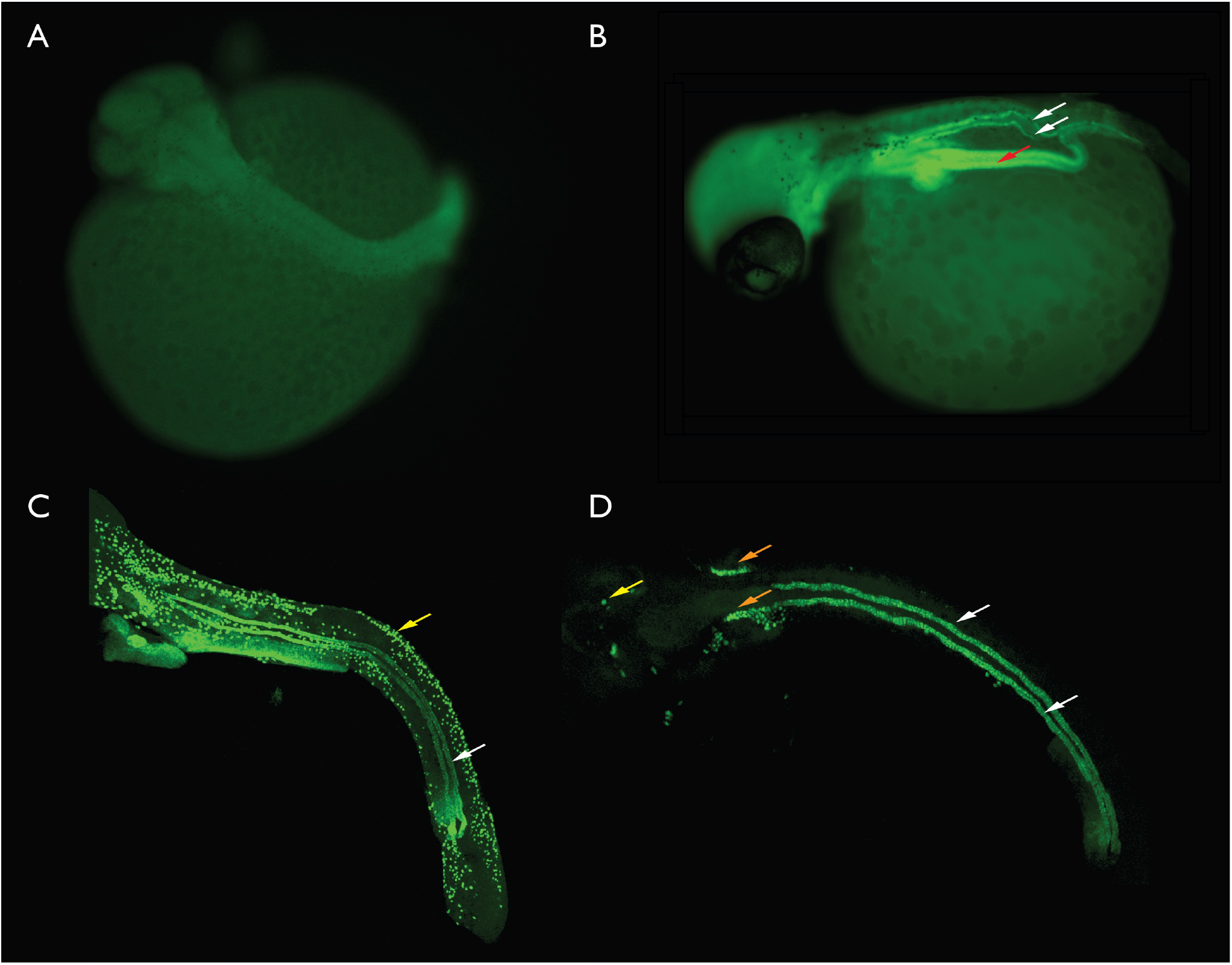
Fluorescent in situ hybridization based labelling of atp1a expression in the developing kidney of bay pipefish embryos. A. Late segmentation stage, no kidney is visible. B. Early pharyngula, paired kidney tubules are present and atp1a positive (white arrows). The gut tube is also fluorescent (red arrow). C. Scanning confocal generated image stack illustrating the kidney tubules (white arrow) and ionocytes (yellow arrow) in a late pharyngula embryo. D. Late pharyngula Scanning confocal generated image stack. This stack was collected only in the plane of the kidney so most other labelled structures were not present. An unknown structure (orange arrows), possibly the developing gills and some ionocytes (yellow arrow) were also present in the image stack.

A range of different embryonic stages, from late segmentation up until late hatching were processed via FISH to visualize the structure of the aglomerular kidney and are illustrated in Figure 5. The nephric system in embryonic pipefishes consists of a pair of epithelial tubules extending anteriorly from the cloaca to just posterior to the head, and located ventral to the somites and notochord. There is an arc in each tubule as the kidney nears the proctodeum, and this probably reflects the kidney bending around the gut tube. The pipefish embryonic kidney looks very similar from pharyngula stages up until hatching. No branching was observed. In order to reduced the number of distracting structures in images, for example the gut, embryos that had undergone FISH were sampled on a confocal microscope. This allowed us to collect a stack of optical sections spanning the kidney, but stopping prior to sampling including other structures. These stacks of optical sections can then be viewed as maximal projections, where all the data is collapsed into a single 2D image. Maximal projections of two different later stage embryos are illustrated in Fig. 5. In the first example some putative ionocytes were also present in the scan (Fig. 5C). In Fig. 5D some scattered ionocytes are present, and another structure, possibly the developing gill system, is also stained. The kidney tubules are clearly visible in both samples.

Scans made using confocal microscopy and the stack of overlapping Z-sections were then used to reconstruct 3D models of the entire kidney. AMIRA software was used to build surface models from confocal data, and thresholding used to remove weaker stained structures. This left 3D surface models of essentially only the paired aglomerular nephrons. Examples of two independent reconstructions are presented in Figure 6, along with maximal projections from the same confocal data sets.

**Figure 6.**
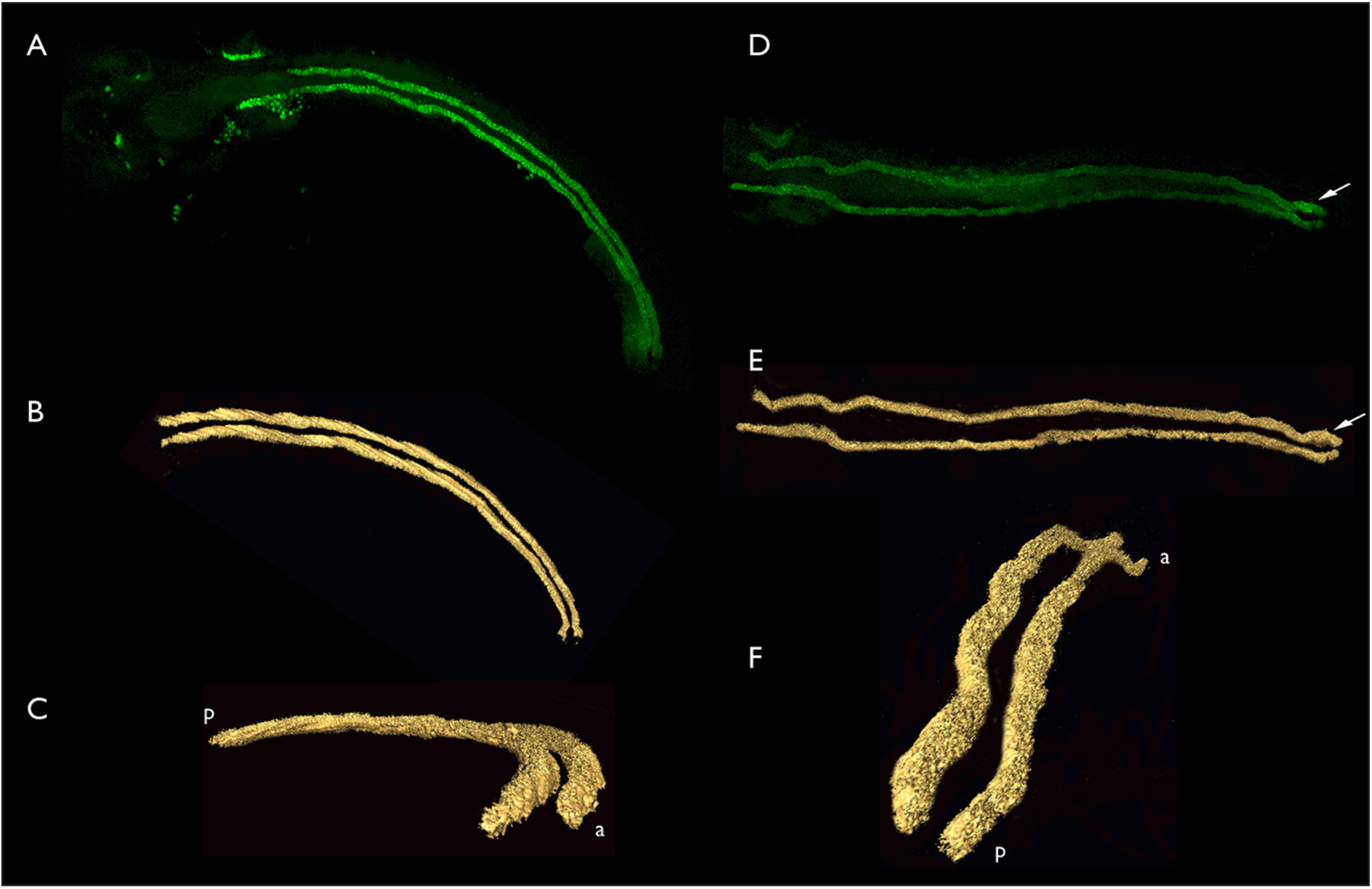
Confocal image stacks and computer generated renderings of aglomerular kidney structure in bay pipefish late pharyngula embryos. A-C, three views of a single set of aglomerular tubules. A. Dorsal view, anterior left, confocal stack. B. AMIRA generated surface model of the data illustrated in panel A. C. The same model as illustrated in panel B rotated so that the anterior portion of the kidney is on the lower right, closest to the viewer. a, anterior; p, posterior. D-F, three views of a single set of aglmoerular tubules. D. Confocal stack. E. AMIRA surface model of the same set of tubules as illustrated in D. F. The model illustrated in E rotated with the posterior end now closest to the viewer. a, anterior; p, posterior.

## Discussion

The embryonic development of a small number of other Sygnathid species has been described and these are referenced and discussed by Sommer et al. (2012). These authors generated a stage series and suggest a small controlled vocabulary that can be applied to all Sygnathids, including pipefishes, seahorses and sea dragons. The stages are not as complete as those illustrated herein for pipefish, including the developmental window in which we characterized aglomerular kidney development in detail, and they also omit the detailed characterization of fin and snout development. The core stages suggested by Sommer et al., (2012) are early embryogenesis, organogenesis, snout formation and new born. These authors suggest that as fin development differs between species it is not useful for a cross-species staging landmark. Within species however, fin development is one of the easiest features to identify and stage, so serves an important role. While the Sommer *et al*. report is an excellent basis for general Syngnathid embryonic staging, additional stages, including the pharyngula stages characterized in detail here, are also necessary to map embryogenesis and provide landmarks for studies of internal organs, such as the developing aglomerular kidneys documented here.

Within pipefish the degree of brood pouch development varies widely from species to species, ranging from simply gluing embryos onto the ventral surface of the male through to highly evolved sealed pouches only slightly less complex than those of their close relatives, the seahorses. In both pipefishes with sealed pouches and seahorses the ionic conditions within the pouch are similar to those in the blood, and very different to the surrounding seawater. The pouch is lined by complex epithelial foldings that are rich in blood vessels (Kornienko, 2001), providing a protected ionic environment and oxygenation. The pouch in some ways represents an evolutionary analog of the mammalian placenta (Azzarello, 1991; Linton and Soloff, 1964). Like the placenta, the pouch also supplies developing embryos with nutrients (Ripley and Foran, 2006) and protects from microbial infection (Melamed et al., 2005). Although seahorse embryos can not survive outside of the pouch (Linton and Soloff, 1964), we have cultured pipefish embryos of a variety of stages for up to 7 days in standard seawater (data not shown) so the dependance of on the pouch is limited to early stages of *S*.*leptorhynchus* development. As discussed above, the translucent pouch makes observing and approximate staging of embryos very straightforward in this species.

While atp1a powers transepithelial ion flow, the question of how the symports and antiports necessary in a glomerular kidney have changed in an aglomerular kidney remains a mystery. The high level of expression of atp1a in the aglomerular kidney indicates that this protein functions in a similar manner in the aglomerular kidney. However, as these kidneys function without utilizing the filtration/resorption system, exactly how this NA/K exchanger is functioning remains to be determined. The Syngnathids have a compact genome not much larger than that of *Fugu* (Vitturi *et al*., 1998) so it will be interesting to determine if many of filtration/resorption genes have been lost through compaction, or what large scale changes have been made to their regulation.

One final question worth considering is whether the evolution of aglomerular kidneys is associated with the evolution of the brood pouch. In elasmobranchs, embryos need to develop a functional kidney in order to survive in the marine environment. Sharks are live-born, while skates and rays develop within a tough salt impermeable egg case. It is possible that the controlled osmotic environment of the Syngnathid brood pouch played a role in the evolution of aglomerular kidneys by allowing embryos to ignore salt and water balance issues until they develop functional aglomerular kidneys.

## Acknowledgments

PDV would like to thank Ramona deGraaf for showing him where and how to collect pipefish around Bamfield Marine Station and the directors, students and staff of BMSC for their hospitality and assistance over many field seasons. Histology of pipefish embryos was kindly performed by Xiaolan Zhou. All fish were collected under permit from BMSC. This research was supported by the Alberta Heritage Foundation for Medical Research.

## Supplemental table

**Table 10.** Optimal permeabilization conditions necessary for successful FISH for each given stage of bay pipefish embryonic development. These conditions include chemical and physical manipulations of the embryos.

| Embryonic stage | Concentration of Proteinase K ($\mu\text{g/mL}$) | Length of permeabilization (minutes) | Temperature of permeabilization ($^{\circ}\text{C}$) | Additional manipulations |
| --- | --- | --- | --- | --- |
| A | 15 | 15 | 25 | Removed from brood pouch |
| B | 15 | 15 | 25 | Removed from brood pouch |
| C | 15 | 15 | 25 | Removed from brood pouch |
| D | 15 | 15 | 25 | Removed from brood pouch |
| E | 15 | 15 | 25 | Removed from brood pouch |
| F | 15 | 15 | 25 | Removed from brood pouch |
| G | 15 | 15 | 25 | Removed from brood pouch |
| H | 15 | 15 | 25 | Removed from brood pouch |
| I | 15 | 15 | 25 | Portion of tail removed |
| J | 30 | 15 | 25 | Portion of tail removed |
| K | 30 | 15 | 25 | Face and half of tail remove |
| L | 125 | 15 | 60 | Face and half of tail remove |
| M | 125 | 15 | 60 | Face and half of tail remove |
| N | 125 | 15 | 60 | Face and half of tail remove |
| O | 250 | 15 | 60 | Face and half of tail remove |
| P | 250 | 15 | 60 | Face and half of tail remove |
| Q | 250 | 15 | 60 | Face and half of tail remove |

